# Tandem RAman Microscopy (TRAM): integration of spontaneous and coherent Raman scattering offering data fusion analysis to improve optical biosensing

**DOI:** 10.1101/2024.04.05.588208

**Authors:** K. Brzozowski, A. Pieczara, A. Nowakowska, W. Korona, B. Orzechowska, J. Firlej, A. Wislocka-Orlowska, M. Baranska

## Abstract

We provide Tandem RAman Microscopy (TRAM), a cutting-edge multimodal microscope that integrates the methods of Stimulated Raman Scattering (SRS), Coherent anti-Stokes Raman Scattering (CARS), and spontaneous (Resonance) Raman Scattering ((R)RS). The device facilitates sequential Continuous Wave (CW) driven RS imaging to collect full spectra from every sample location and rapid pulsed-wave-driven SRS-CARS scanning at specific wavenumbers, offering a reliable and efficient analytical tool. The fingerprint spectral region can be included in the spectral imaging capabilities of CARS and SRS. Data collected from a sample area using several techniques can be integrated and analyzed, significantly increasing reliability and predictions. We analyzed the *in vitro* model of nonadherent leukocytes (LC) to illustrate the capabilities of this unique system, emphasizing the benefits of measuring the same sample with three different Raman techniques without having to transfer it between microscopes. Data fusion allowed for the correct classification of two subtypes of LC based on the partial least squares (PLS) discrimination, increasing the prediction accuracy from approximately 83% in the case of textural and morphological data (SRS) to 100% when combined with spectral data (SRS and RS). We also present RRS images of LC labeled with astaxanthin, and reference data from SRS and CARS microscopy. Additionally, polystyrene beads were investigated as a non-biological material. The advantages of each Raman technique are utilized when (R)RS, SRS, and CARS are combined into a single device. This paves the way for dependable chemical characterization in a wide range of scientific and industrial fields.

## Introduction

Understanding material structure and composition is essential to deciphering the language of nature and comprehending its behaviors, properties, and functions. However, because of the inherent intricacy and limitations of individual analytical tools, getting detailed information can be difficult.

An analytical method needs to meet several criteria to be deemed reliable and measurement requirements include (i) high accuracy and precision, (ii) high selectivity and sensitivity, (iii) assessment of both qualitative and quantitative properties, (iv) versatility enabling application to a variety of sample types, (v) minimal sample preparation and non-destructiveness, (vi) high speed and high throughput, (vii) robust performance, (viii) easy to use and cost-effective, and (ix) offering high spatial resolution imaging capabilities.^1–5^

The majority of these requirements are met by spectroscopic techniques, including Raman microscopy, and therefore they are widely used in many fields as tools to assess chemical and structural information even from fragile samples.

Raman spectroscopy is a fundamental analytical method in the field of material sciences and biology, offering non-invasive probing of the molecular composition of samples. It is based on Raman effect,^1^ a phenomenon in which incident light is inelastically scattered by a sample. The so-called Raman shift provides information about the molecular vibrational energy of sample components, which allows for their detailed chemical analysis.

Among Raman techniques, spontaneous Raman Scattering (RS) spectroscopy, which records the linear response of a sample to the applied electric field, is the most widely used. Various applications of RS in chemistry,^2^ material science,^3^ biology,^4^ and medicine^5^ are focused on the identification and quantification of substances, the study of molecular interactions, and the monitoring of biochemical processes in live cells. Despite the universality and comprehensiveness of the RS technique, it has some limitations, particularly in terms of speed and sensitivity,^6^ which is an obstacle when dealing with biological samples that may be susceptible to damage under prolonged exposure to highly focused laser light. However, the sensitivity can be significantly enhanced by taking advantage of the electronic transition, as is the case for resonance RS (RRS). Regarding speed, data acquisition in RS is typically conducted using translation stages equipped with piezo-elements, which provide high accuracy down to nanometers. Although such imaging seems to be relatively slow, it is not the translation speed of the stage that limits it, but the time required to collect the Raman spectrum at a given measurement point. Therefore, in most cases, there is no need to use faster scanners in RS, because the speed limitations inherent to RS result from its linear nature and small cross-section of the phenomena, which requires longer exposure times at each measurement point than, for example, in the case of fluorescence. Moreover, a major problem for RS is fluorescence, which often obscures the Raman signal, making molecular composition detection difficult.^7^

Coherent anti-Stokes Raman Scattering (CARS) and Stimulated Raman Scattering (SRS) overcome the intrinsic challenges encountered in RS by providing enhanced signal strength and rapid measurements. Both techniques pioneered Coherent Raman Scattering (CRS), since their invention revolutionized vibrational imaging with a video-rate timescale.^8,9^ For example, in the case of biological material, the RS signal for a single pixel is collected over several hundred milliseconds, whereas detection of the CARS or SRS signals takes tens of microseconds. Therefore, in imaging using nonlinear optics techniques, where information about one component is obtained at the expense of the entire Raman spectrum, the measurement time is several orders of magnitude shorter than in the case of RS.^10^ That is achieved by employing galvo-scanners, which enable measurements by the movement of a laser beam as an alternative to piezo-stage scanning.^11^

The fundamentals of CARS are based on a four-wave mixing process encompassed by joint interaction between coherently incident Stokes and pump photons with the sample to generate a new field at anti-Stokes frequency.^10^ Current applications of CARS exhibit growing interest in the fields of biomedical samples endoscopy^11^ and intracellular metabolism^8,12^ while offering an enhanced Raman signal with an increased selectivity of imaged molecule vibrations. The capability of tuning the pump and Stokes frequency difference in CARS to match the symmetric stretch vibrations of the CH_2_ groups showing a large Raman scattering cross section has been demonstrated in multiple studies on lipid metabolism in cells,^13^ tissues,^14^ and model organisms.^15^ Despite multiple advantages, the CARS signal suffers from the contribution of a non-resonant background, requiring the application of e.g. phase retrieval algorithms to obtain valuable information.^8,9^

Similarly to CARS, SRS spectroscopy is based on the 3rd order nonlinear effect that enhances the Raman signal through the interaction of the sample with a pump beam and a Stokes beam. Compared to CARS, SRS’s imaging capabilities are based on rapid galvo-scanners and picosecond lasers, enable ultrafast imaging of specific Raman bands and better discrimination against non-resonant background.^16^ SRS has proven to be particularly beneficial for biological applications, as high-throughput measurements allow the capturing of dynamic processes^16^ and minimize damage^17^ to the sample.

Using CARS and SRS imaging, two nonlinear optics techniques that have a distinct physical origin from RS phenomena, offers benefits and therefore provides a more comprehensive insight into the vibrational state of samples.^18^ However, despite the advantages of CARS and SRS over RS, there are a few challenges and considerations with CRS techniques that need to be taken into account. The undeniable advantage of CRS is its high confocality and the possibility to register a discrete signal that comes from the vibration of a certain energy. On the other hand, analysis based solely on a single selected signal may be difficult when the energy of vibrations from various molecules is similar and their Raman bands overlap. A reliable Raman microscope should therefore combine the advantages of the speed and efficiency of nonlinear optic spectroscopic techniques with the robustness of spectroscopic interpretation through the analysis of the Raman spectrum obtained with RS. Therefore, we propose the Tandem RAman Microscope (TRAM), which integrates the best of the Raman scattering techniques, i.e. (R)RS, CARS, and SRS, into a single platform.

Because CARS and SRS signals are generated simultaneously at different wavelengths when the sample is illuminated with pump and Stokes beams, it is possible to detect both CARS and SRS at the same time using a single microscope.^19^ This approach has been applied effectively to study the dynamics of biological systems.^20,21^

Some attempts have already been made to combine CARS or/and SRS with RS. The combination of CARS and RS was used for a quantitative analysis of lipid bodies,^22^ and revealed differences in the level of unsaturation and the abundance of exogenous fatty acids in adipocytes. Both CARS and RS measurements were performed on the same platform with the use of a picosecond laser and galvo-scanner. However, this approach was further enriched by independently conducted analysis of lipid droplets through SRS and RS, on separate samples, different from those measured by CARS and RS. That experimental setup offered the capability for simultaneous RS-SRS, and RS-CARS measurements, but with certain constraints, primarily the use of a picosecond excitation source with no flexibility for choosing a UV-VIS laser source for RS excitation, especially useful for RRS. A combined analysis using SRS and RS imaging has also been demonstrated to distinguish normal tissue from cancerous tissue in mouse fibrosarcoma.^23^ RS showed the potential for improved accuracy diagnostics for future spectral histopathology. Both imaging techniques were also used to study the distribution of lipids in the skin.^24^ The combined analysis helped to reduce the dominant protein background and cross-sensitivity for mixed Raman bands, allowing the extraction of spectra of skin lipids. In the pharmaceutical area, integration of SRS and RS was used to follow the spatial distribution of amlodipine besylate in tablets, which is a commonly used blood pressure-lowering drug.^25^ Both techniques have been applied to the cancer detection and drug development process.^26^ However, in this case, SRS and RS measurements were performed using separate microscopes, making it difficult to maintain the same conditions and measurement area.

RS and SRS analysis also have the potential for real-time monitoring of dynamic processes. The capability of SRS to capture dynamic changes in molecular composition has already been demonstrated in the progression of degenerative diseases, such as atherosclerosis^27^ or lipogenesis.^28^ Moreover, imaging sensitivity and selectivity can be further enhanced by labeling, i.e. utilizing specific Raman reporters containing an isotopic or alkyne tag.^27^ In a study of atherosclerosis, Stiebing et al. showed repeated measurements of the same cells over time to track their metabolism in real-time using both SRS and RS, but separately. The importance of tracking changes over time in the same cells to capture cell-to-cell variability has been demonstrated, rather than taking measurements at different times of different cellular populations at corresponding intervals.^27^ Therefore, the ability to measure the same cells over time and with different techniques is crucial to fully understand the complexity of the systems under study, as well as to correlate the information obtained by different methods and minimize the impact of cell mobility and the need for spatial calibration.

In material science, Ploetz et al.^29^ utilized non-invasive optical microscopy and spectroscopy techniques for the *in situ* correlative analysis of metal-organic framework particles. They demonstrated that the crystal shape of MIL-88A significantly influences its optical absorption and explored the uniformity of the water distribution in MOF-801, a material promising for the harvesting of water from humid air. Imaging techniques, such as RS and CARS, alongside other methods like four-wave mixing, sum frequency generation, second harmonic generation, two-photon excitation, and fluorescence were utilized there. The complementarity of RS and SRS measurements is also used for material characterization and design, allowing *in situ* studies on dynamic processes such as ion transport^30^ or polymerization.^31^

Despite previous efforts, no technological solution has been presented that combines the three Raman modalities, (R)RS, CARS, and SRS, in one device as effectively as our tandem system. TRAM integrates these modalities, co-located on a single platform, enabling both spectral characterization and imaging. Key advantages of TRAM include (i) the use of a continuous wave (CW) UV-VIS laser, which allows for flexible measurement of (R)RS without the constraints typically associated with picosecond lasers used for CRS, and (ii) the inclusion of two scanning methods compatible with (R)RS and CRS measurements, i.e. a high-precision piezo-stage and an ultra-fast galvo-scanner.

## Methods

### Tandem RAman Microscopy (TRAM)

We used TRAM for in-depth spectroscopic analysis using a multimodal approach that combines (R)RS, SRS, and CARS.

The integration of the *Alpha 300Ri* spectrometer (*WITec* GmbH, Germany) equipped with a CCD detector (*Andor Technology* Ltd., Northern Ireland) and a 600 grooves/mm grating (BLZ = 500 nm) in conjunction with electronics and *Control SIX 6.1* software (*WITec* GmbH, Germany) was used for RS microscopy, together with a 532 nm CW laser (*WITec* GmbH, Germany) with an average power output of 50 mW. The same electronics and software were used for SRS and CARS, which makes the system exceptionally useful.

The core of the SRS and CARS setup in the TRAM is the picosecond *Lazurite* laser system (*Fluence* sp. z o.o., Poland). It includes a fiber-based Stokes laser that can produce an average power of 450 mW at a 1029 nm wavelength. Additionally, the system features a built-in Optical Parametric Oscillator (OPO) that generates a tunable pump beam, ranging from 750 to 950 nm, with an average power of 100 mW. Both the Stokes and pump beams produce synchronized pulses of 2 ps with a repetition rate of 20 MHz. To get the beams ready for CRS imaging, they are first overlapped in space using a dichroic mirror and aligned in time with an optical delay line (ODL) in the path of the pump beam. The Stokes beam is modulated at a 4 MHz frequency using an Acousto-Optic Modulator (AOM).

The beams are directed to the sample using a dual-axis piezo-mirror galvo-scanner attached to a *Nikon Ti2* inverted microscope (*Nikon Corp.*, Japan). An *UPLXAPO40X* NA 0.95 40× (*Olympus Corp.*, Japan) air objective lens focuses the laser beams onto the sample. For signal collection, we use the same objective lens (RS epi-detection), and an *MRD07620* 50× NA 1.0 (*Nikon Corp.*, Japan) water dipping objective lens (SRS and CARS detection in transmission). An *SM1PD1* photodiode (*Thorlabs Inc*., US) and *SR865A* lock-in amplifier (*Stanford Research Systems*, US), which demodulates the signal, are used for SRS collection, while the *R9110* photomultiplier tube (*Hamamatsu Photonics*, Japan) plus the *SR570* preamplifier (*Stanford Research Systems*, CA, US) are used for CARS collection.

For the CARS collection, we avoid non-resonant background by applying optical filters that remove Rayleigh and SHG signals. Additionally, Stokes and pump signals were carefully removed by additional band-pass and low-pass filters to not disturb the CARS signal. **Fig. 1a–b**. presents the TRAM setup along with energy diagrams of the methods.

Additional information regarding the influence of SRS/CARS pixel integration time on image quality and a comparison between scanning methods, specifically galvo and piezo scanners in RS, can be found in the Supplementary Materials. These aspects are thoroughly discussed and presented there.

### Experimental Parameters

RS measurements of cells were performed using TRAM set to approx. 24 mW and 12 mW average power of the 532 nm beam for label-free and labeled cells, respectively, a pixel dwell time (PDT) of 500 ms, and a fixed pixel size of 1 µm x 1 µm. Frame size was set to 40 µm x 40 µm for label-free HL-60 live cell measurements, 50 µm x 40 µm for labeled HL-60 fixed cell measurements, and 22 µm x 32 µm for label-free HL-60 and Jurkat cell line mixture for testing the model based on the fusion of RS and SRS data. Also, polystyrene beads (PB) were examined. In the case of PS beads, the power was set to approx. 12 mW, PDT of 50 ms, fixed pixel size of 1 µm x 1 µm, and a frame size of 50 µm x 60 µm.

For CRS imaging of cells, the system was configured to an average power of approx. 24 mW of the Stokes beam and approx. 12 mW of the pump beam. CRS frames for label-free and labeled HL-60 measurements were captured with a PDT of 900 µs, a fixed pixel size of 500 nm x 500 nm, and frame sizes of 40 µm x 40 µm, and 50 µm x 40 µm respectively. The SRS frames for measurements of the mixture of HL-60 and Jurkat cell lines were captured with a PDT of 900 µs, a fixed pixel size of 333 nm x 333 nm, and a frame size of 100 µm x 100 µm. In the case of PS beads, the power was set to approx. 6 mW of the Stokes beam and 3 mW of the pump beam, PDT of 900 µs, fixed pixel size of 500 nm x 500 nm, and a frame size of 50 µm x 60 µm.

Our TRAM setup offers two modes of sample scanning. Scanning using a piezo-stage is utilized only in (R)RS measurements, while scanning using a galvo-scanner is employed in SRS and CARS, and also in (R)RS measurements.

### Cells preparation

Human promyelocytic leukemia HL-60 (ECACC 98070106) cells and T-cell acute lymphoblastic leukemia (T-ALL) Jurkat (ECACC 88042803) cells were cultured in Roswell Park Memorial Institute (RPMI) 1640 medium supplemented with glutamine, 10% Fetal Bovine Serum (FBS) and 1% antibiotic (penicili, streptomycin, and amphotericin). Cell cultures were kept at 37 ^°^ C, 5% CO_2_/95% in an air-humidified incubator. The cell density was maintained below one million cells per milliliter. HL-60 cells were divided into three groups: one group was incubated with 200 µM of palmitic acid (Sigma Aldrich) for 24h; the second group was incubated with 10 µM of astaxanthin (AXT) (Sigma Aldrich) for 6 h; and the third group consisted of control cells. Live cell measurements were conducted on the first group and part of the control sample. Just before the measurements, the cells were rinsed with warm PBS. The remaining control HL-60 cells and the AXT sample were measured fixed, as well as Jurkat cells. HL-60 and Jurkat cells were first washed with warm PBS and then fixed with 0.5% glutaraldehyde for 10 min, then washed three times with PBS. To build a model based on the fusion of RS and SRS data, fixed HL-60 and Jurkat cells were measured separately and in a mixture of cells, prepared right before data acquisition in a proportion of 50% : 50%. Before each measurement, cells were placed onto CaF_2_ slides, and suspended in a droplet. Measurements were performed after cell sedimentation and immobilization. The fixed samples were preserved in PBS buffer at a temperature of 4 ° C until Raman measurements were performed.

### Image processing and software

Data from RS, SRS, and CARS were analyzed by employing *Project SIX 6.1 software* from *WITec*. Preprocessing of Raman spectra included Cosmic Ray Removal (CRR; filter size: 4; dynamic factor: 4) and Background Subtraction (BG; polynomial order: 3), then a KMC (*k*- means cluster) analysis was performed. All measurements and initial data analysis were operated by one software, *Project SIX 6.1*. Origin 2021 (OriginLab Corporation) was used for data presentation. To build a predictive model based on the fusion of RS and SRS, the dataset was analyzed using Python scripts, prepared in-house. Preprocessing of RS spectra included CRR (spectrapepper Python package, cosmicdd, th: 10, asy: 0.6745, m: 10), Singular Value Decomposition (SVD), denoising (numpy Python package, 3 SVs), extended multiplicative signal correction (EMSC, biospectools Python package), and Sensitive Non-linear Iterative Peak clipping (SNIP) baseline correction (iterations: 100, smoothing window: 11). The segmentation of the RS maps and extraction of the mean spectra of individual cells was performed automatically on the basis of the calculation of the Pearson correlation coefficients with the reference cellular spectrum. RS and CRS image segmentation was done using the scikit-image Python package (watershed and canny edge detector algorithms). For RS spectra, Principal Component Analysis (PCA) was performed for the dimensionality reduction using the scikit-learn Python package (7 PCs, explaining in total 95.83% of data variability). Before segmentation, SRS images were corrected by subtracting the background signals collected at 2800 cm^-1^. For single SRS images of cells, a ratiometric analysis (2850 cm^−1^/2930 cm^−1^) was performed. To extract useful features for analysis of segmented CRS ratiometric images of single cells, the Grey-Level Cooccurrence Matrix (GLCM) algorithm, algorithm (scikit-image Python package), and textural parameters such as energy, correlation, dissimilarity, homogeneity, contrast, and Shannon entropy were calculated. Furthermore, the cell size was calculated as a morphological parameter. Partial Least-Squares Discriminant Analysis (PLS-DA) was used to obtain classification models with the PLS Toolbox with MIA Toolbox 9.3.1 software (Eigenvector Research, Inc., US). The analysis was performed separately on the RS spectra, on values of the textural and morphological parameters, and on the merged spectral, textural, and morphological data. After combining spectral, textural, and morphological datasets, data standardization was performed. For each model, 70% of the data was used for the calibration of each model (with cross-validation of venetian blinds) and 30% of the data was used for the validation of the model.

## Results

We demonstrate the versatility of the TRAM setup by examining its applications in the fields of biological and material sciences. The challenge in biological research is, among others, the observation of subcellular composition, organelle activity, and the dynamics of cell metabolism. Materials science is a diverse and interdisciplinary field that focuses on the development and synthesis of new substances, including metals, polymers, semiconductors, and ceramics. In both research areas, the correlation between structure and functionality is crucial. Spectroscopy has also found its niche in this field, showing the use of SRS ^11–14,32–36^ and CARS ^15–17,37–42^ to study the properties of various materials.

### TRAM in biological sciences: subcellular analysis

Here we successfully employed TRAM to assess the metabolic state of live leukemia cells (LC) simultaneously using three spectroscopic modalities, i.e. RS, SRS, and CARS, over a wide spectral range under the same conditions (**Fig. 2a-c**). RS was used to register the full spectra and select the bands for subsequent SRS and CARS measurements. SRS and CARS images (**Fig*. 3*a-b**) were collected using a pump beam at: 1/ 791 nm (2930 cm^−1^), mainly corresponding to proteins, to show cell morphology, 2/ 796 nm (2850 cm^−1^), mainly corresponding to lipids, and 3/ 879 nm (1660 cm^-1^), corresponding to lipids and proteins. RS images (**Fig*. 3*c**) were obtained by integrating the characteristic Raman bands and provide information on cell morphology (2835-3025 cm^−1^, CH stretching), the subcellular localization of endogenous lipids (2845–2855 cm^-1^, ν(CH) in CH_2_ stretching), and subcellular localization of lipids and proteins (1630-^1^690 cm^-1^, v(C=C) stretching, amide I). Moreover, the *K*-Mean Cluster (KMC) analysis showed the spatial distribution of the subcellular components (**Fig*. 3*d**). The mean spectra of unsaturated lipids, saturated lipids, nucleus, cytoplasm, and membrane are presented in **Fig*. 3*e**. A strong correlation is observed between the SRS and RS images that depict mainly the morphology of lipid droplets. The discrepancy between the SRS and CARS images, notably at 2930 cm□¹, is attributed to the variation in the focal planes of cells across different frame locations combined with a weak CARS signal.

**Figure. 2.**
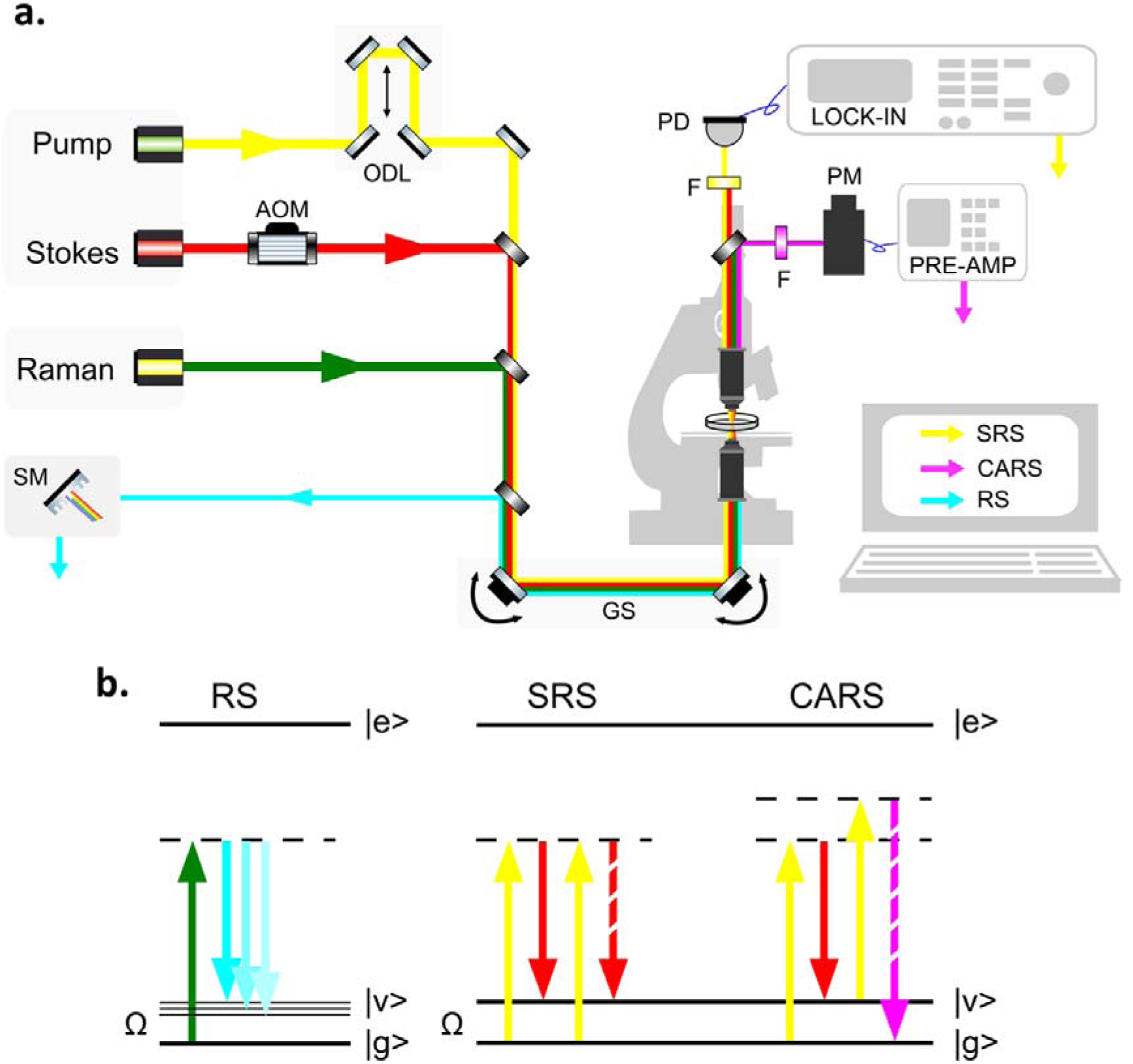
**a.** TRAM setup; AOM – Acousto-Optic Modulator, ODL – Optical Delay Line, F – optical filters, PD – photodiode, PM – photomultiplier tube, GS – galvo-scanner, SM - spectrometer. **b**. Energy-level diagrams of RS, SRS, and CARS.

**Figure. 3.**
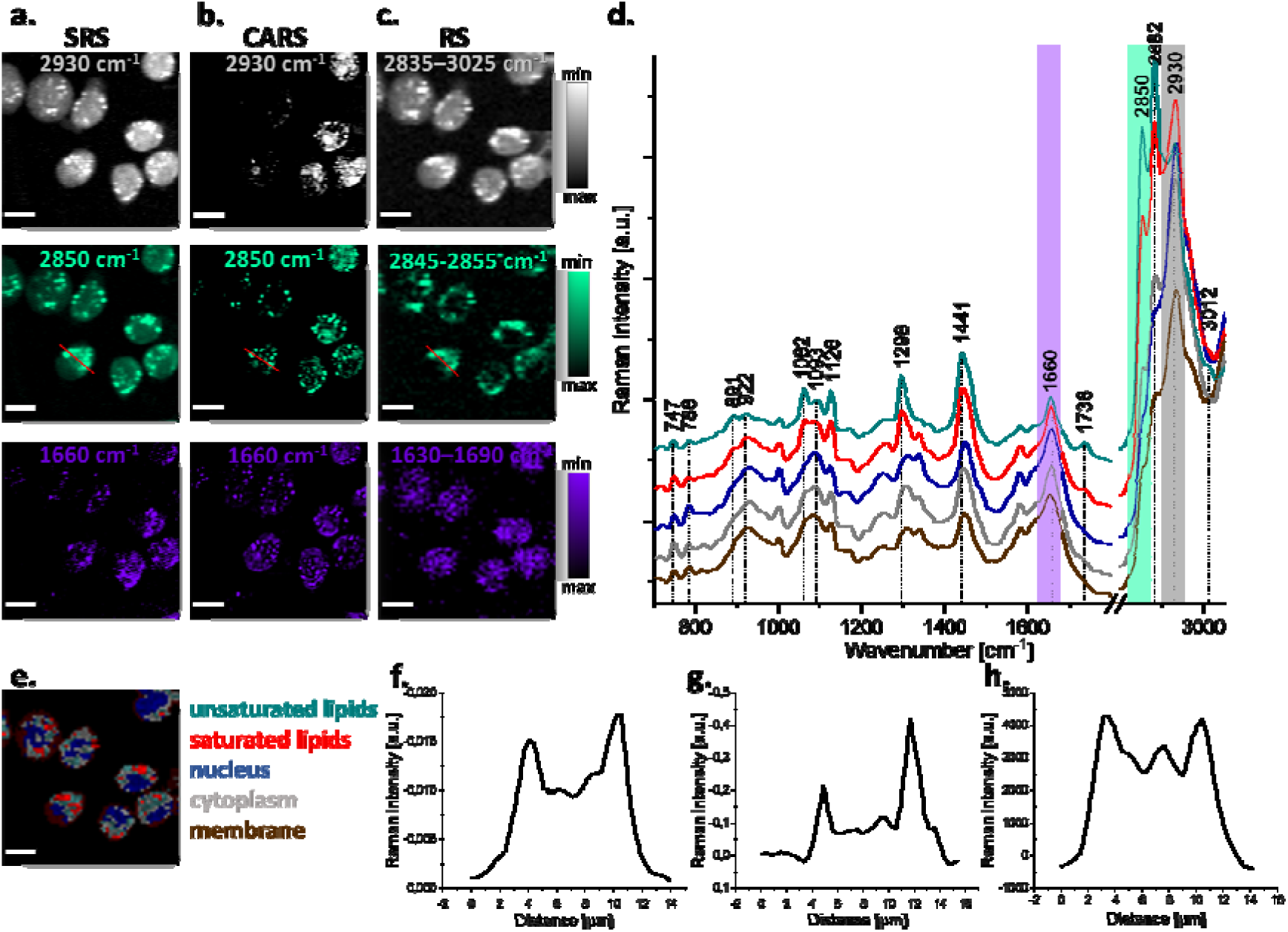
SRS/CARS/RS images of live HL-60 cells. **a**. – **b.** SRS and CARS images were collected using a pump beam of 791 nm (2930 cm^−1^, organic matter), 795 nm (2850 cm^−1^, lipids), and 879 nm (1660 cm^-1^, lipids and proteins) respectively. **c**. Representative RS images were obtained by the integration of the selected Raman bands at 2835–3025 cm^−1^ (organic matter), 2845–2855 cm^−1^ (lipids), and 1630–1690 cm^-1^ (lipids and proteins), respectively. **d**. KMC map, showing the distribution of cell components in HL-60 cells. **e.** Averaged Raman spectra of cell components: unsaturated lipids (green), saturated lipids (red), nucleus (blue), cytoplasm (grey), and membrane (brown). **f–h**. Intensity profiles of Raman signals collected along the cross-section in images showing the distribution of lipids (highlighted in red) obtained using SRS, CARS, and RS, respectively. Scale bar: 10µm. SRS and RS scans were performed using a galvo-scanner and a piezo-stage, respectively.

**Table 1.**
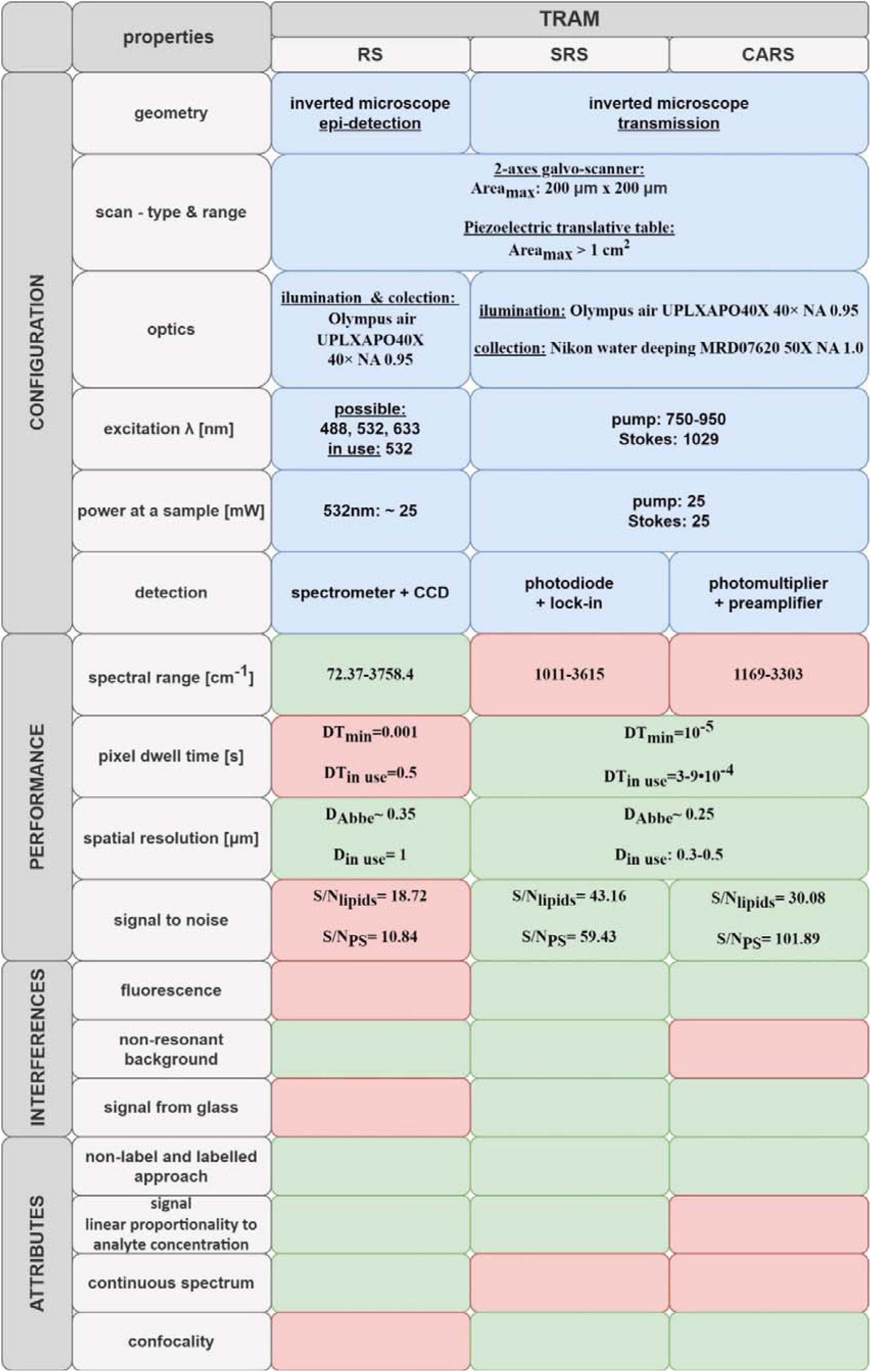
Comparison of technical parameters for RS, SRS, CARS, and TRAM.

The images of SRS, CARS, and RS show slightly different lipid distribution. This disparity arises from the relatively broad integration range employed to visualize the heterogeneous lipid fraction in RS (10 cm□¹), juxtaposed with the discrete specific wavenumber (2850 cm□¹) chosen for the SRS and CARS measurements.

The measurement time for RS and CRS was significantly different, i.e. the imaging of cells using the RS took several minutes (∼ 25 min), while the SRS and CARS images were obtained in a few seconds (14 s), although the step size was twice as small for SRS and CARS (0.5 µm) in comparison to RS (1 µm). This shows the advantage of CRS imaging in terms of speed.

The SRS images are sharp, and the contrast between the signal and the background is high. The signal-to-noise ratio (S/N) ) **(Fig*. 3*f-h**), calculated along the cross section in images showing the distribution of lipids, is around two and half times higher for SRS images (43.16) and about two times higher for CARS images (30.08) in comparison to the corresponding RS images (18.72). The SRS signal does not suffer from a non-resonant background, which greatly improves the quality of the SRS images and lowers the noise. Additionally, the measurement step size is two times smaller for SRS and CARS than for RS, ensuring better visualization of details in the cell structure for the first approach, e.g. the perinuclear area was visualized distinctly using the SRS technique.

Resonance-enhanced RS enables a very sensitive detection of a specific group of chemical components. To meet the requirements to obtain RRS it is necessary to adjust the energy of the excitation wavelength to the electronic transitions of the molecule ^43^. In TRAM we incorporated a CW 532 nm laser, which gives the possibility of RRS measurements. For the first time, we demonstrate RRS images together with CRS data obtained on the same microscope on the example of cytochromes in the mitochondria of live leukemic cells (**Fig. 3**) ^44^. SRS images (***Fig. 3*3a**) were collected using a pump beam at 879 nm (1660 cm^−1^), corresponding to lipids and proteins and showing cell morphology, and 885 nm (1583 cm^-1^), mainly corresponding to cytochromes c and b ^45^. RS images of representative cells (***Fig. 3*3b**) were obtained by integrating the characteristic Raman bands for lipids and proteins (1630-1690 cm^-1^, v(C=C) stretching, amide I) and heme proteins (1573-1593 cm^-1^, v(C C_m_) stretching of the heme group) ^46^. Using KMC analysis, it was possible to show the spatial distribution and mean spectra of the subcellular components (***Fig. 3*3c, 3d**), including mitochondria. The Raman bands of the cytochrome are strong and well resolved (752, 1128, 1312, 1583 cm^-1^)^45,47,48^. Consequently, the distribution of cytochrome can be easily tracked using the 1583 cm^-1^ band (**Fig. 3b**). On the contrary, the SRS image at 1583 cm^-1^ (**Fig. 3a**) does not show mitochondria as clearly as the RRS image. To avoid photodamage, SRS and RS imaging was performed on different sets of cells.

Alongside live cells in a label-free approach, we also measured fixed HL-60 cells labelled with AXT. AXT is a naturally occurring compound that belongs to the carotenoid family. This red color pigment has antioxidant and anti-inflammatory properties that are attributed to its distinctive molecular structure, which enables it to effectively eliminate free radicals and reactive oxygen species ^49,50^. AXT, when excited by a 532 nm laser, exhibits a significant resonance-enhanced Raman signal. The most characteristic bands of AXT are 1520 cm^−1^, 1157 cm^-1^, and 1005 cm^-1^. SRS images were collected for the 1520 cm^-1^ band characteristic for AXT and the 1660 cm^-1^ band corresponding to proteins and lipids, while RRS images show the spatial distribution of AXT (1500-1540 cm^-1^) and cellular morphology (organic matter: 2830-3030 cm^-1^, and lipids and proteins: 1575-1705 cm^-1^). Both RS and SRS data clearly show AXT localized inside the labelled cells compared to the control sample (**Fig. 4**). What is more, RRS spectra show a significant enhancement of the AXT signal, exhibiting the intense bands specific to AXT, which significantly facilitates the detection of this compound in cells with high sensitivity.

**Figure. 4.**
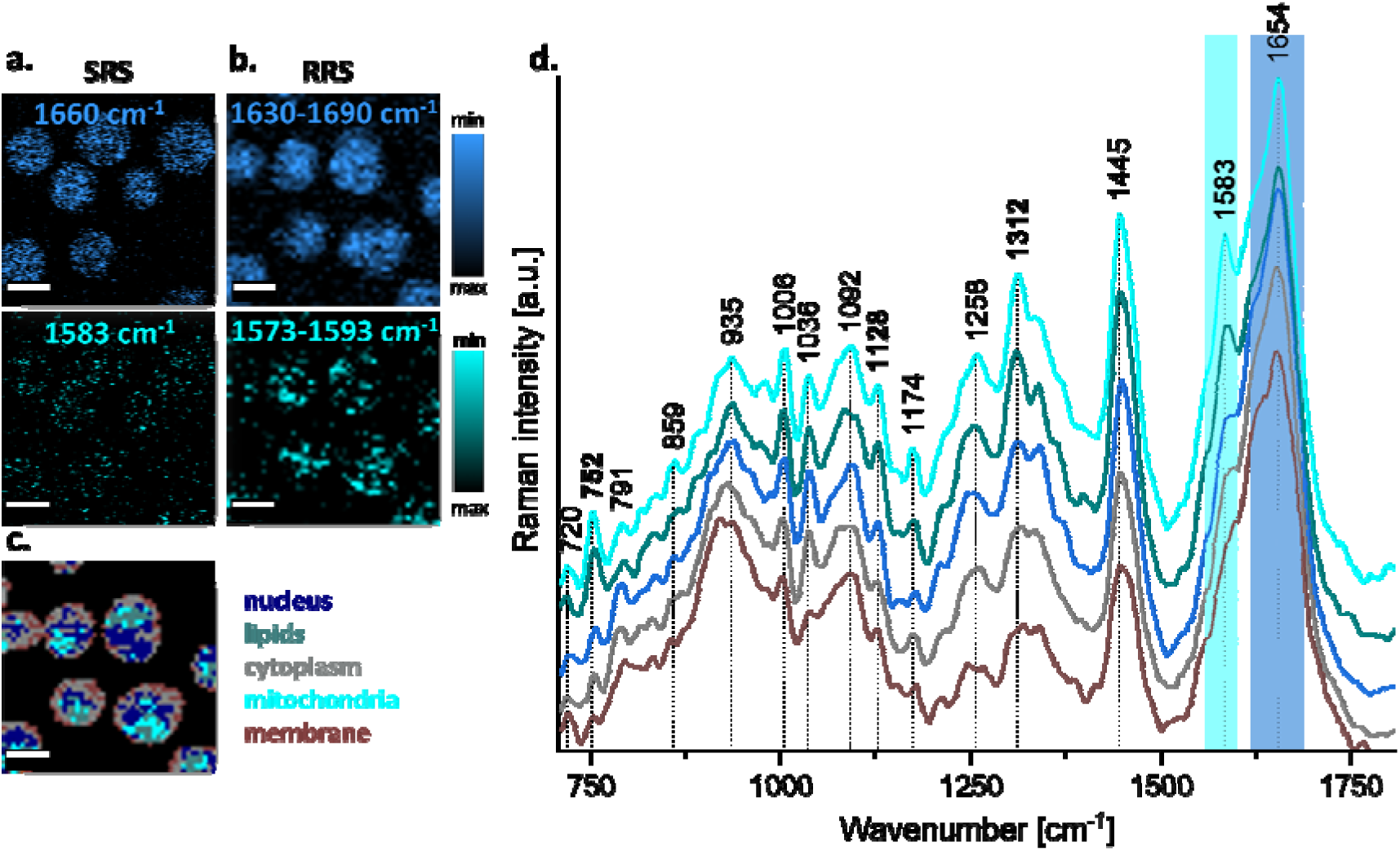
SRS/RS images of live HL-60 cells. **a**. SRS images were collected using a pump beam of 879 nm (1660 cm^−1^, lipids and proteins), and 885 nm (1583 cm^−1^, mainly cytochromes in mitochondria), respectively. **b.** Representative RS images were obtained by integration of the selected Raman bands at 1630-1690 cm^-1^ (lipids and proteins), and 1573-1593cm^−1^ (mainly cytochromes in mitochondria, exhibiting RRS), respectively. **c**. KMC map, showing the distribution of cell components in HL-60 cells. **d.** Averaged Raman spectra of cell components: lipids (green), mitochondria (light blue), nucleus (blue), cytoplasm (grey), and membrane (brown). Scale bar: 10µm. SRS and RS scans were performed using a galvo-scanner.

**Figure. 4a.**
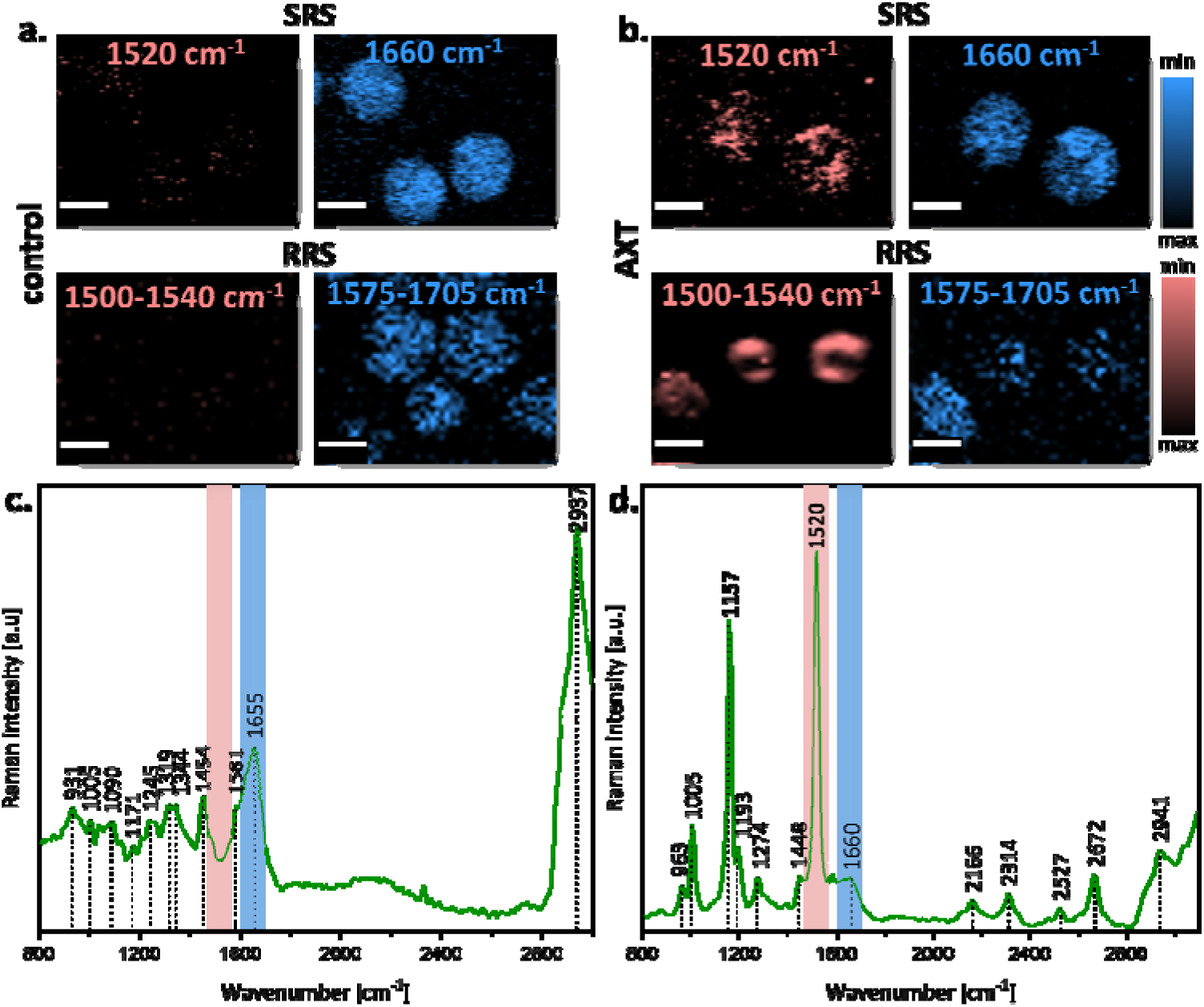
SRS/RS images of live HL-60 cells. **a**. Control sample. **b.** AXN labelled sample; SRS images were collected using a pump beam of 879 nm (1660 cm^−1^, lipids and proteins), and 890 nm (1520 cm^−1^, AXN Raman band). RRS images were obtained by the integration of the selected Raman bands at 2830-3030 cm^-1^ (organic matter), 1575-1705 cm^−1^ (lipids and proteins), and 1500-1540 cm^-1^ (AXN Raman band) **c.** Raman spectrum of control cells. **d.** Raman spectrum of AXT in labelled cells. Scale bar: 10µm. SRS and RS scans were performed using a galvo-scanner.

### TRAM in biological sciences: data fusion to improve cell phenotyping

An important aspect of biomedical analysis is cell phenotyping, which allows for reliable identification of subpopulations of cells in the context of their state and function. Based on the single spectroscopic method, it is possible to build classification models to be used for the prediction of the cell subtype ^51–56^. With TRAM, there is a unique possibility to combine data collected from various spectroscopic methods and perform data fusion analysis to improve the performance of the models and extend their applicability to more complex problems, including the identification of populations of cells in highly heterogeneous neoplasms, such as leukemia ^57,58^.

We performed RS and SRS imaging of *in vitro* models of two types of LC, i.e. Jurkat cell line, representing T-cell Acute Lymphoblastic Leukemia (T-ALL), and HL-60, representing acute promyelocytic leukemia (APL). To enable subsequent data fusion, RS spectra and SRS maps were recorded for exactly the same cells, which, using TRAM, is very easy as it does not require moving the sample. We collected a series of SRS maps, using a pump beam at: 1/ 791 nm (2930 cm^−1^), mainly corresponding to proteins, 2/ 795 nm (2850 cm^−1^), mainly corresponding to lipids, and 3/ 800 nm (2800 cm^-1^) as background signals for further correction of artefacts (**Fig. 5a**). Data was collected from control Jurkat and HL-60 cells, and from a mixture of T-ALL and APL to test the performance of the classification models (cells were mixed in a proportion of 50%:50%). Segmentation of the RS and SRS images was performed automatically using edge detector algorithms (**Fig. 5a and Fig. 5b**). For RS data, mean spectra of cells were extracted, and to reduce the dimensionality of the dataset, PCA was used. For single SRS images of cells, a ratiometric analysis was performed (2850 cm^−1^ / 2930 cm^−1^), simultaneously including information on the chemical and spatial structure of lipids and proteins (**Fig. 5a**). To extract the features for all SRS ratiometric images, the GLCM algorithm was used, and the textural parameters, such as energy, correlation, dissimilarity, homogeneity, contrast, and Shannon entropy, were calculated ^58^. Furthermore, the cell size was calculated as a morphological parameter. For the fused data of SRS and RS of LC subtypes (T-ALL *vs* APL) we built the PLS-DA models. Classification models were obtained separately on features determined from SRS images (**Fig. 5a**), on RS spectra (**Fig. 5b**) and on combined RS and SRS data (**Fig. 5c**). 70% of the data was used for the calibration of each model (with cross-validation of venetian blinds, Cal) and 30% of the data were used for the validation of the model (Val). The model trained on values of the textural and morphological parameters of the ratiometric SRS images gave an accuracy equal to 70.83 % (Cal) and 83.33 % (Val) (**Fig. 5d**). An analogous model, but built only on spectral data, gave accuracy equal to 88.46 % (Cal) and 100% (Val) (**Fig. 5d**). However, by joining spectral, morphological, and textural information, improved accuracy of the models was achieved, i.e. up to 96.15% (Cal), and 100% (Val) (**Fig. 5d**).

**Figure. 5.**
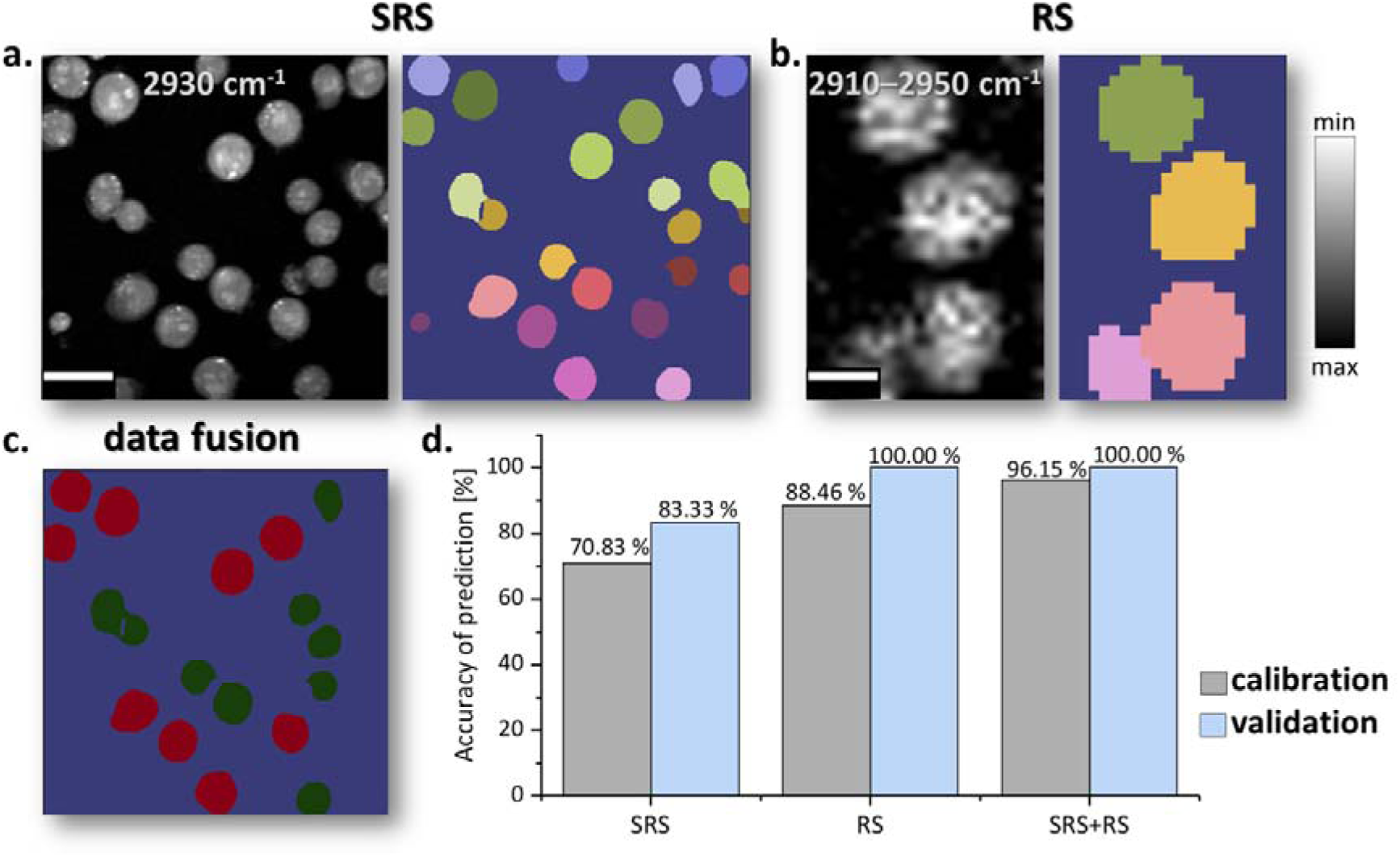
Data fusion for improved LC subtyping. **a.** Analysis of SRS images; SRS images were collected using a pump beam of 791 nm (2930 cm^−1^) and 795 nm (2850 cm^−1^). Based on the SRS maps registered mainly for proteins at 2930 cm^−1^, characterizing cell morphology, image segmentation was done and ratiometric analysis was performed for the SRS maps of single cells (2850 cm^−1^/2930 cm^−1^). **b.** Analysis of RS spectra; For the same cells for which SRS measurements were performed, RS spectra were recorded using a 532 nm CW laser. The RS maps were segmented based on the Raman bands that are primarily specific for proteins (2910 – 2950 cm^−1^) and the RS spectra of individual cells were extracted. **c.** PLS-DA model to classify APL (red) and T-ALL (green) subtypes of leukemia based on the fusion of data from SRS ratiometric images and RS spectra. The model obtained was used to predict subtypes of LC in a mixture of cells (50% T-ALL (green):50% APL (red)). **d.** Accuracy of prediction calculated for calibration and validation data for classification models obtained separately on features determined from SRS images (SRS), on RS spectra (RS), and combined RS and SRS data (SRS + RS).

The data fusion model was further used to classify the cells in the mixture (**Fig 5c)**. The algorithm predicted cells in a 50%:50% mix, equal to the initial proportion of APL and T-ALL cells in the sample.

This example demonstrates the potential of TRAM in the context of data fusion for an improved diagnosis of LC ^58^. Since RS and SRS data were recorded for the same cells, it was possible to perform data fusion (RS + SRS) and improve the efficiency of classification of the T-ALL and PML types of LC based only on SRS images. This approach opens possibilities for the construction of efficient prediction models that can be used in clinics for more complex real-life situations.

### TRAM in material science

To present the versatility of our setup, the CRS/RS measurements of the PS beads were done (**Fig. 6**). We compared the signal-to-noise ratio calculated along the cross-section in images showing the distribution of the 3055 cm^-1^ band for SRS, CARS, and RS. In the case of CRS, this ratio is about six and ten times higher, respectively, for CARS (101.89) and SRS (59.43) images compared to the corresponding RS (10.84) image. The SRS signal does not suffer from a non-resonant background, which highly improves the quality of SRS maps, and lowers the noise. However, for the 2905 cm□¹ band, the weaker CARS signal appears to be excessively strong, suggesting potential additional effects imposed on CARS observation. Conversely, for the 1604 cm□¹ band, the CARS signal seems to be significantly stronger than expected, surpassing even the SRS signal strength, which may be indicative of potential additional non-resonant background.

**Figure. 6.**
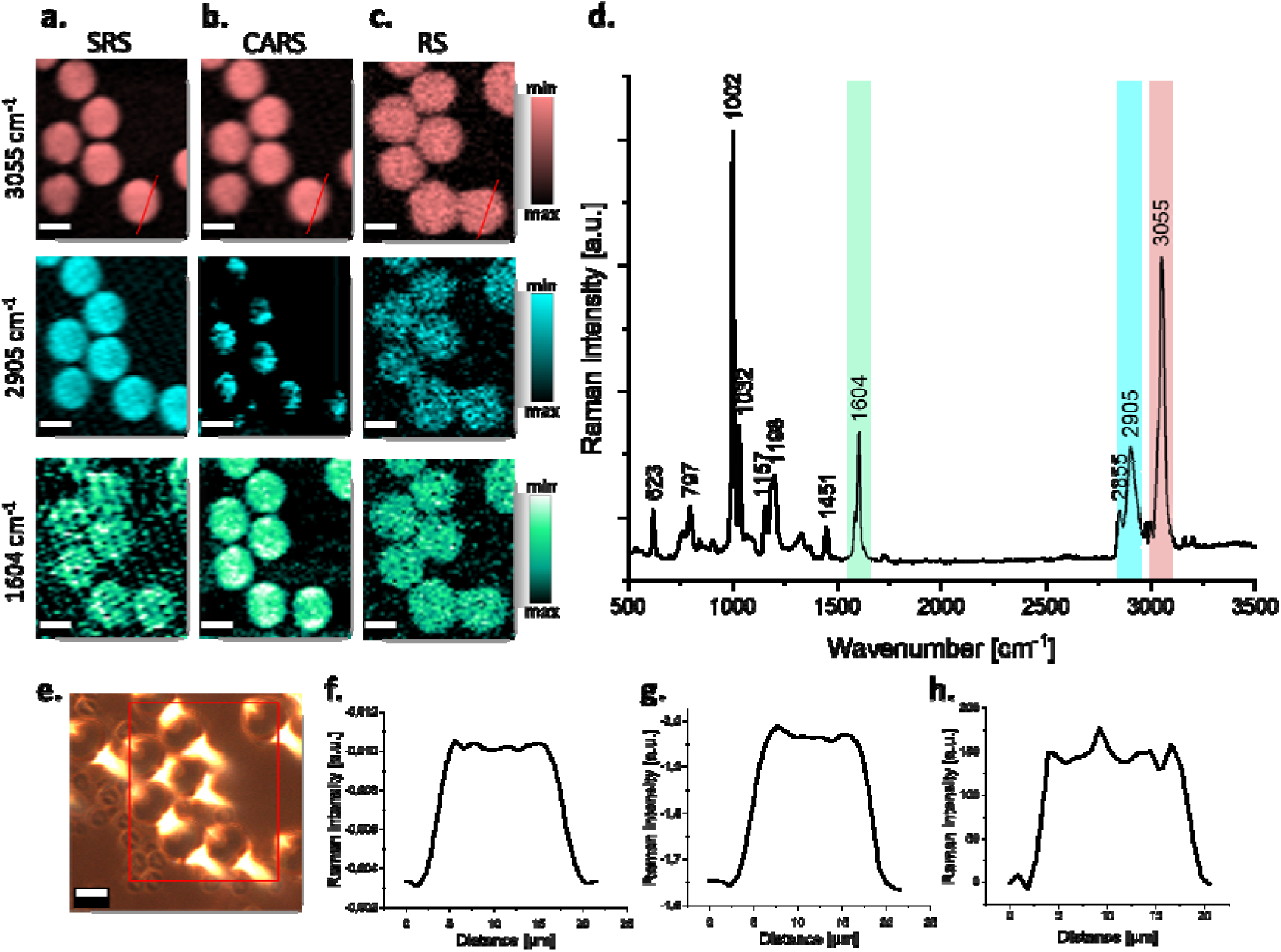
SRS/CARS/RS images of PS beads. **a. – b.** SRS and CARS images were collected using a pump beam of 783 nm (3055 cm^−1^), 792 nm (2905 cm^−1^), and 883 nm (1604 cm^-1^), respectively. **c**. Representative RS images were obtained by the integration of the selected Raman bands at 3055 cm^−1^, 2905 cm^−1^, and 1604 cm^-1^ respectively. **d**. Bright field of the measured field. **e.** The averaged Raman spectra of PS beads. **f-h**. Intensity profiles of Raman signal collected along the cross-section (highlighted in red) using SRS, CARS, and RS, respectively. Scale bar: 10µm. SRS and RS scans were performed respectively using a galvo-scanner and a piezo-stage.

## Discussion

The integration of (R)RS, CARS, and SRS into a single multimodal microscope represents a significant step forward in the field of spectroscopy and microscopy. This innovative combination harnesses the strengths of these techniques, offering a comprehensive tool for the detailed analysis, especially of biological samples including live cells.

To sum up, the advantages and implications of TRAM are the following:

### 1. Multimodal approach

Simultaneous usage of both (R)RS and CRS techniques in a single device is rare. While each technique offers valuable insights, its combination as a single microscope amplifies its advantages. This multimodal approach allows for the simultaneous collection of both broad spectral information and the targeted analysis of specific Raman bands.

### 2. CRS combined with CW (R)RS

Integrating CRS techniques with CW (R)RS in a single microscopy system offers significant advantages. Primarily, the use of a CW laser for (R)RS measurements, as an alternative to the picosecond pulsed lasers typically employed for SRS and CARS, can be particularly beneficial. The CW laser facilitates the use of UV-VIS wavelengths, which are essential when enhancing resonance Raman effects for certain biomolecules. We demonstrated the efficiency of this approach in RRS analysis of cytochrome in a label-free way and for the AXT-labeled sample. Moreover, CW lasers present a lower risk of sample damage compared to high-energy pulsed lasers. This is crucial in (R)RS, where prolonged illumination at a single spot is necessary due to the inherently weak Raman signals, requiring slow scan speeds. In contrast, SRS and CARS benefit from the short, intense illumination of pulsed lasers that quickly capture images, minimizing potential photodamage. Additionally, the use of a CW laser in (R)RS allows for better control over the illumination intensity, further reducing the risk of thermal damage to sensitive samples. This combination not only extends the capabilities of Raman microscopy by accommodating a broader range of experimental conditions and sample types but also enhances data quality and safety during prolonged imaging sessions.

### 3. Two scanning modes

The ability to scan samples using two scanning options, i.e. a piezo-stage and galvo-scanner, significantly enhances the versatility and efficiency of the system. The use of the piezo-stage for scanning in RS measurements allows for a thorough analysis of large sample areas, which is crucial to obtain representative and comprehensive spectroscopic results. On the other hand, employing galvo-scanner in the CRS techniques enables much faster imaging of specific molecular components, i.e. while the piezo-stage allows for achieving PDT of the order of ms, the galvo-scanner allows for PDT of μs. TRAM is particularly useful in studies that require high resolution and speed, such as real-time imaging of dynamic biological processes. Combining both scanning techniques in one device opens new possibilities for investigating the structure and dynamics of complex samples while significantly reducing the time needed to conduct experiments. This innovative configuration not only facilitates more accurate investigations, but also contributes to the development of more advanced spectroscopic analytic methods.

Using an objective lens with glass correction to avoid glass signal overlapping with the Raman spectrum of the sample, galvo-scan for RS is possible. It can be particularly useful for samples with a large Raman cross-section which can be scanned rapidly and TRAM may change the perception of (R)RS microscopy as a relatively slow method.

### 4. Consistency in measurements

One of the advantages of this integrated approach is the ability to multimodally measure the same sample under the same conditions. This is particularly important for non-adhesive cells, such as suspension cells not attached to a dish. The system enables direct data integration by correlating RS, SRS, and CARS signals at the single-pixel level. A major benefit of a tandem system is that once the sample is placed, it can be measured using different techniques without any displacement, ensuring that the area under investigation is not changed. This capability is invaluable, particularly for studies involving sensitive or dynamic cellular processes, as it allows uninterrupted, consistent observation, and further comprehensive analysis including data fusion. By minimizing the handling of the sample and reducing exposure to potentially harmful conditions, the accuracy and reliability of the collected data are maintained.

### 5. Enhanced efficiency by data fusion

Integration of the CRS and (R)RS techniques into a single device significantly increases the efficiency of the analysis. The speed of the CRS techniques, which allows for ultra-fast, high-quality imaging and molecular characterization perfectly complements the RS method that provides full spectral information. This complementarity enables the comprehensive utilization of both techniques within one microscope, not only shortening the time required to perform CRS and RS measurements but also increasing the precision and reliability of the collected data.

Data fusion analysis can significantly improve the performance of the classification models. In the context of biological samples, this approach can improve cell phenotyping, and hence diagnostic capabilities. It is possible to integrate spectral data from RS with morphological and textural features of CRS. In the example presented here, this dual-data approach allowed us to refine our classification models and use PLS-DA to achieve remarkable improvements to prediction accuracy, i.e. from 83.33 % with textural and morphological data alone (CRS) to 100 % when combined with spectral data (CRS+RS). The results underscore the potential of TRAM in leveraging data fusion for enhanced biomedical applications, affirming the synergy between CRS imaging and spectroscopy in a unified analytical platform.

### 6. Further upgrade

TRAM offers the potential for expansion to include additional imaging techniques such as the Second Harmonic Generation (SHG). By employing appropriate filters, dichroic mirrors, and additional detection, it is possible to split the signal for simultaneous detection of CARS, SRS, and SHG signals. Such an integration would allow concurrent imaging of multiple properties of samples with unprecedented accuracy and detail. While incorporation of CARS provides vibrational contrast, adding another layer of information to the images generated by SRS, the SHG utilizing the generation of the second harmonic, would enable the observation of structures like collagen or muscle fibers with unique contrast.^30,31^ Separation of signals and simultaneous detection of these signals open new avenues in the structural and functional studies of various cell types and tissues, offering comprehensive insights into their complex interactions and dynamics.

## Conclusions

TRAM integrates CARS, SRS, and (R)RS within one microscope and is a significant step forward in the field of spectroscopy and microscopy. This opens up new opportunities in research and facilitates sample analysis, while setting new high standards in biology, chemistry, medicine, and materials science. The unique capabilities of the device can be further improved and developed.

Even though it is not yet cost-effective and portable, TRAM offers a methodology that enables the comprehensive observation of Raman phenomena across diverse physical origins, encompassing both linear and nonlinear effects. Hence, this methodology does not require the reference techniques in advanced optical imaging, while ensuring complex sample characterization.

Despite these innovations, there are still challenges associated with the sensitivity and specificity of Raman microscopy. However, the integration of CARS, SRS, and (R)RS within a single microscope facilitates addressing these issues, enabling more effective and precise adjustments. As a result, the parameters of the measurement can be better controlled, compared, optimized, and data from several techniques can be integrated for the further improvement of the sensitivity and specificity of spectral imaging, and prediction of classification models.

