## Supplemental Files for "Tandem RAman Microscopy (TRAM): integration of spontaneous and coherent Raman scattering offering data fusion analysis to improve optical biosensing"

**Supplementary Information**

**SRS/CARS pixel integration time influence on image quality**

For imaging of biological material using our TRAM, the most challenging samples are endothelial cells (EC) (or in general, adherent cells). This is because adherent cells spread out on the surface of the substrate and then reach thicknesses of the order of 10-30 µm ^1,2^. Compared to blood cells, which are round and are of the order of 10-20 µm in size (or more for macrophages, for example)^3^, the SRS and CARS signal is less intense for EC compared to leukocytes (LC). In the case of RS, for which measurements are made on glass, the contribution of signals from the substrate is also greater for adherent cells in comparison to suspension LC due to the smaller thickness of adherent cells. Therefore, we have tested how the imaging time relates to the quality of the maps obtained for EC. For this purpose, normal primary Human Aortic Endothelial Cells (HAEC) incubated with palmitic acid (PA) were investigated with SRS and CARS microscopy.

**Cells preparation**

HAEC were cultured in EGM™-2 Endothelial Cell Growth Medium-2 (EGM™-2), supplemented with 5 % heat-inactivated [fetal bovine serum](https://www.sciencedirect.com/topics/medicine-and-dentistry/fetal-bovine-serum) (FBS), 0.04 % Hydrocortisone, 0.4 % hFGF-B (human fibroblastic growth factor), 0.1 % VEGF (vascular endothelial growth factor, 0.1 % R3-IGF-1 (insulin-like- growth factor), 0.1 % Ascorbic Acid, 0.1 % hEGF (epidermal growth factor), 0,1 % GA-1000 (antibiotic mixture) and 0.1 % Heparin. Cells were grown in glass bottom dishes at 37 °C in 98 % air / 5 % CO_2_ atmosphere. Cells were seeded directly onto glass bottom dishes 24h before the incubation at 37 °C in 98 % air / 5 % CO_2_ atmosphere, this allowed the cells sufficient time to grow and spread, reaching an appropriate confluence. Then cells were incubated with 200 µM of PA (Sigma Aldrich) for 24h. PA was first saponified in NaOH. Then heated to 70 ° C to liquefy it. To facilitate the transport of PA through the cell membrane, saponified PA was conjugated with bovine serum albumin (BSA). Following incubation, cells were gently rinsed with PBS and fixed for five minutes at room temperature with 2.5% glutaraldehyde. The fixed cells were then maintained in Dulbecco`s Phosphate Buffered Saline (DPBS) and kept at 4°C constant temperature until measurements were taken. Just before measurements, the cells were rinsed with warm PBS.

**Cells imaging**

The SRS/CARS images were taken using the TRAM setup. The setup was tuned to 2850 cm^-1^. The average power of 24 mW of Stokes 1029 nm and 12 mW of pump 795 nm were used. The image size was 40 μm x 40 μm. The scan resolution was 200 px x 200 px. Integration time *vs.* picture quality is provided in Fig. S1 and Fig. S2 for SRS and CARS, respectively.

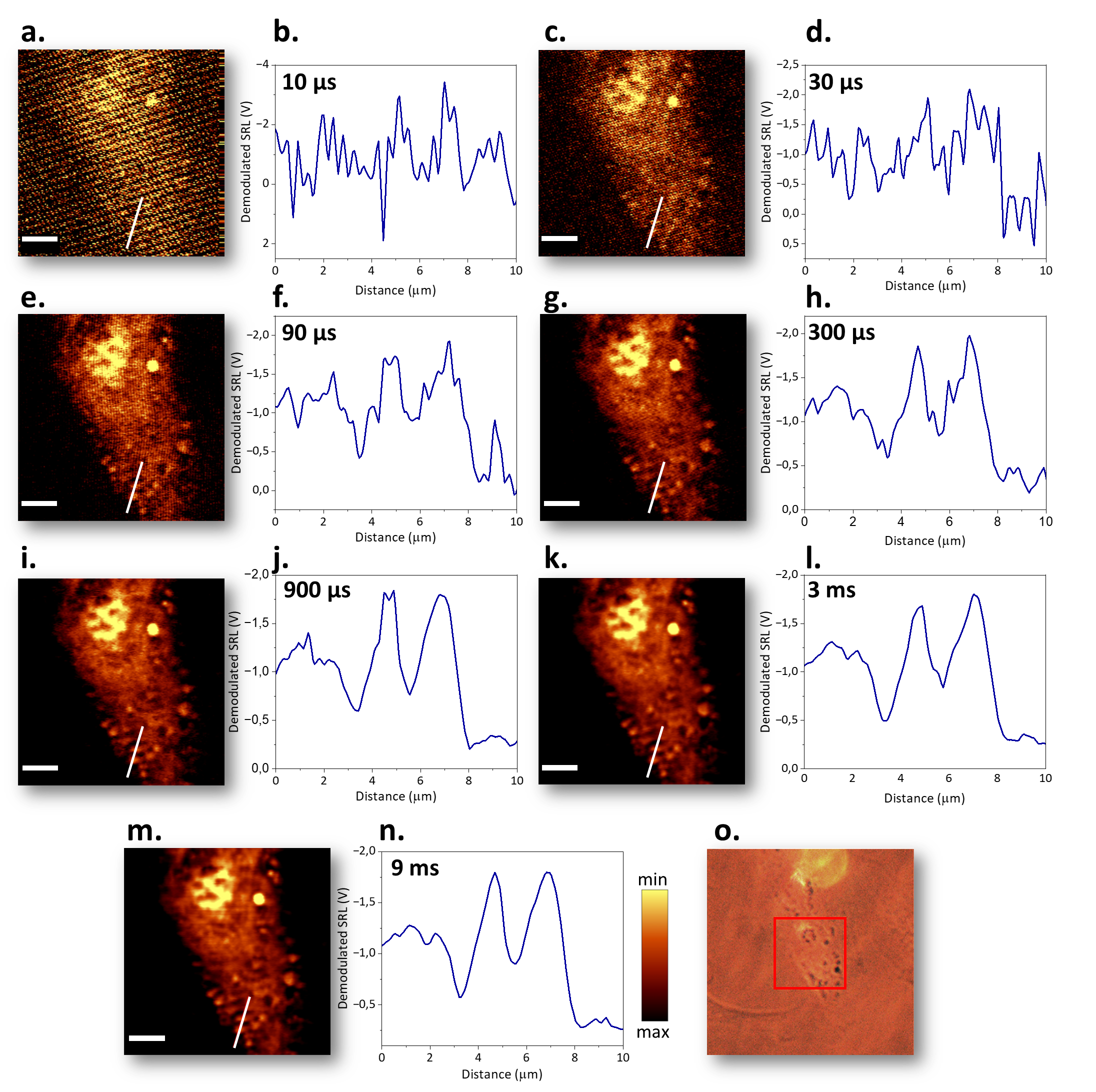

***Fig. S1 SRS pixel integration time influence on image quality.*** *SRS images of HAEC cells incubated with PA were collected using different integration times:* ***a.*** *10 μs,* ***c.*** *30 μs,* ***e.*** *90 μs,* ***g.*** *300 μs,* ***i.*** *900 μs,* ***k.*** *3 ms,* ***g.*** *9 ms.* *Intensity profiles of Raman signals collected using different integration times:* ***b.*** *10 μs,* ***d.*** *30 μs,* ***f.*** *90 μs,* ***h.*** *300 μs,* ***j.*** *900 μs,* ***l.*** *3 ms,* ***n.*** *9 ms along the cross-section (highlighted in white) in images showing the distribution of lipids obtained for SRS.* ***o.*** *Optical preview of the sample with marked 40 μm x 40 μm imaging area. Scale bar: 7 µm.*

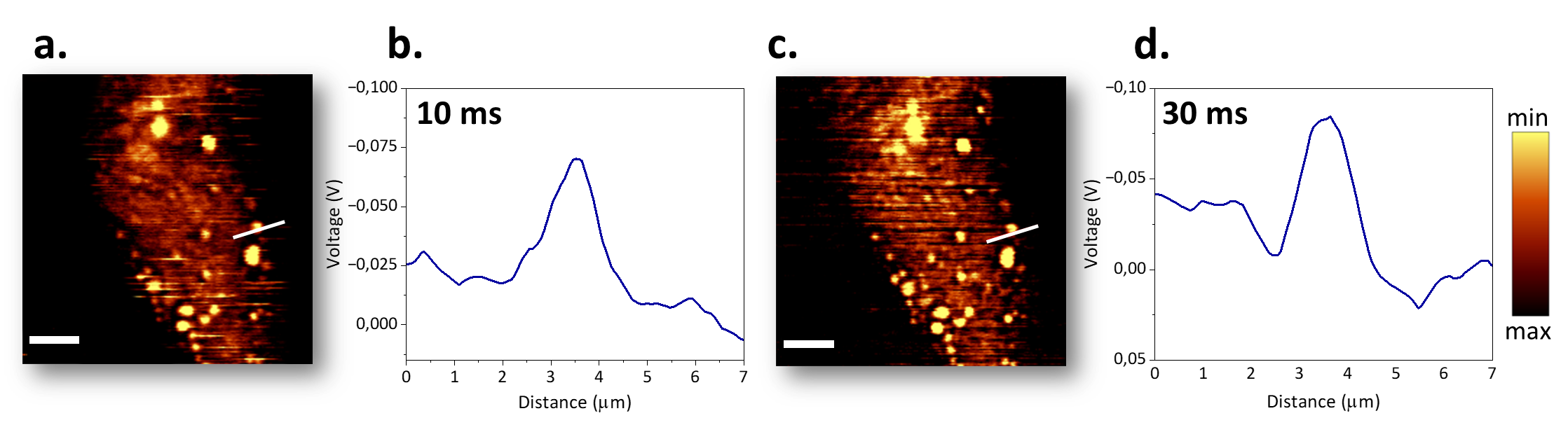

***Fig. S2 CARS pixel integration time influence on image quality.*** *SRS images of HAEC cells incubated with PA were collected using different integration times:* ***a.*** *10 μs,* ***c.*** *30 μs.* *Intensity profiles of Raman signals collected using different integration times:* ***b.*** *10 μs,* ***d.*** *30 μs along the cross-section (highlighted in white) in images showing the distribution of lipids obtained for CARS. Scale bar: 7 µm.*

Taking into account speed vs quality, optimal pixel dwell time for SRS imaging using TRAM is 900 μs. Further integration time increase doesn’t significantly improve image quality while elongating storage time meaningly.

The CARS signal is collected simultaneously with SRS and satisfactory images emerge even at a minimal pixel dwell time of 10 μs. However, all structures identifiable by CARS become apparent at a dwell time of 30 μs dwell time, and extending this duration further does not enhance the results. Additionally, due to the concurrent storage of SRS and CARS frames, we utilize the same integration time for both methods. It is worth noting that the structures visible using CARS slightly differ from those observed with SRS. Those differences can be attributed to aspects such as contrast mechanisms, molecular sensitivity, background influence, and spatial and spectral resolution that differ for both methods.

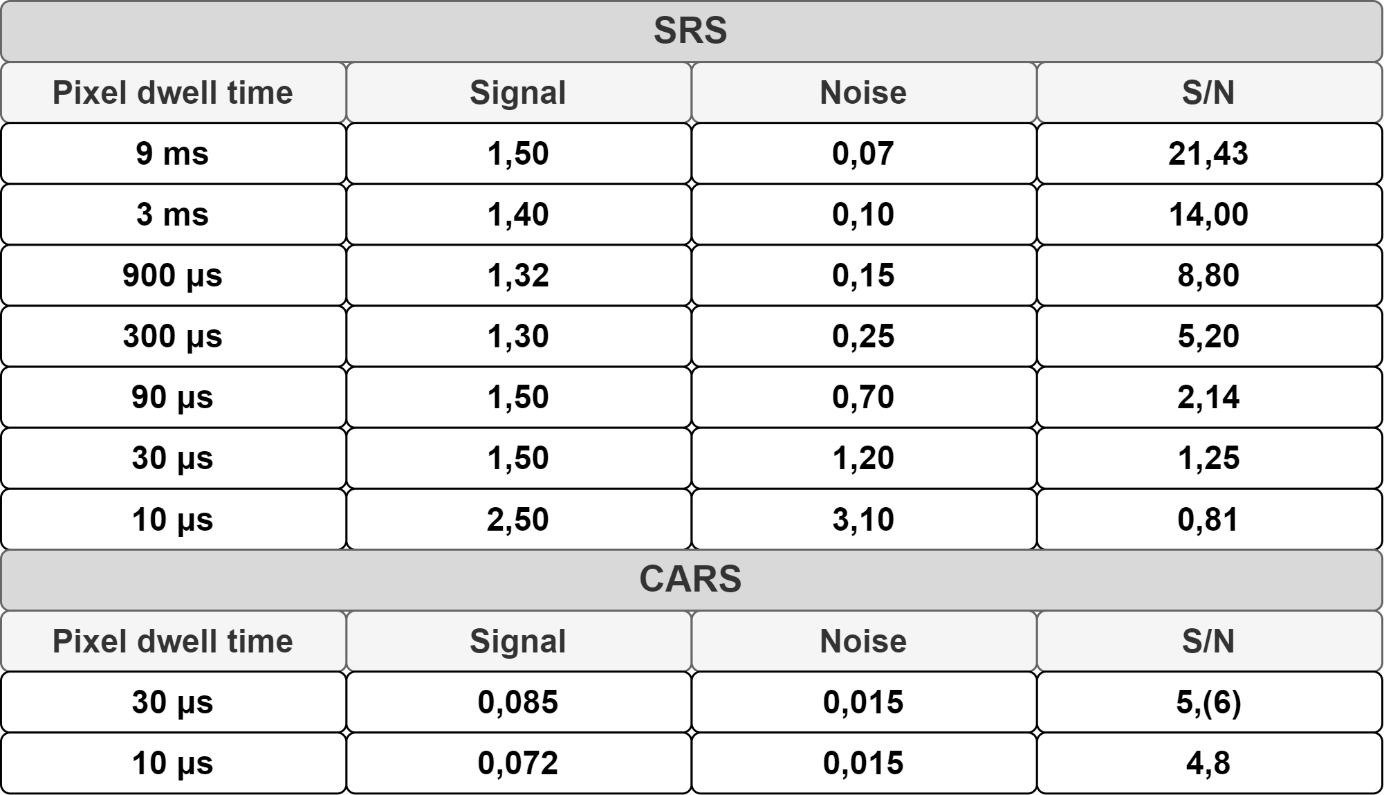

***Table S1. S/N for SRS and CARS***

**Comparison between scanning methods – galvo and piezo scanner in RS**

Our TRAM setup allows scanning using a piezo-stage and a galvo-mirror scanner. For CARS and SRS, we use the galvo due to its speed, while for RS, the piezo-stage suffices in terms of speed. We have compared both scanning methods during RS measurements of Acute Promyelocytic Leukemia cells (HL-60).

**Cells preparation**

HL-60 cells were cultured in (RPMI), supplemented with 10 % heat-inactivated fetal bovine serum (FBS) and 1 % antibiotic (mixed of streptomycin and penicillin). Cells were grown in six-well plate at 37 °C in 98 % air / 5% CO_2_ atmosphere. Cells were cultured for 24 h before measurements. After being removed from the incubator, the cells were centrifuged and washed with warm DPBS solution tree times. After that cells were placed on the microscopic stage for imaging.

**Cells imaging**

Images were taken using TRAM setup with RS module. 532nm laser beam was used for spectra acquisition. 25 mW average power was used. Image size was 33μm x 33μm. The scan resolution was 33 px x 33 px. Integration time was 0.5 s. RS images were obtained by integrating spectra in the range of 2845–3045 cm^-1^ (characteristic of organic matter).

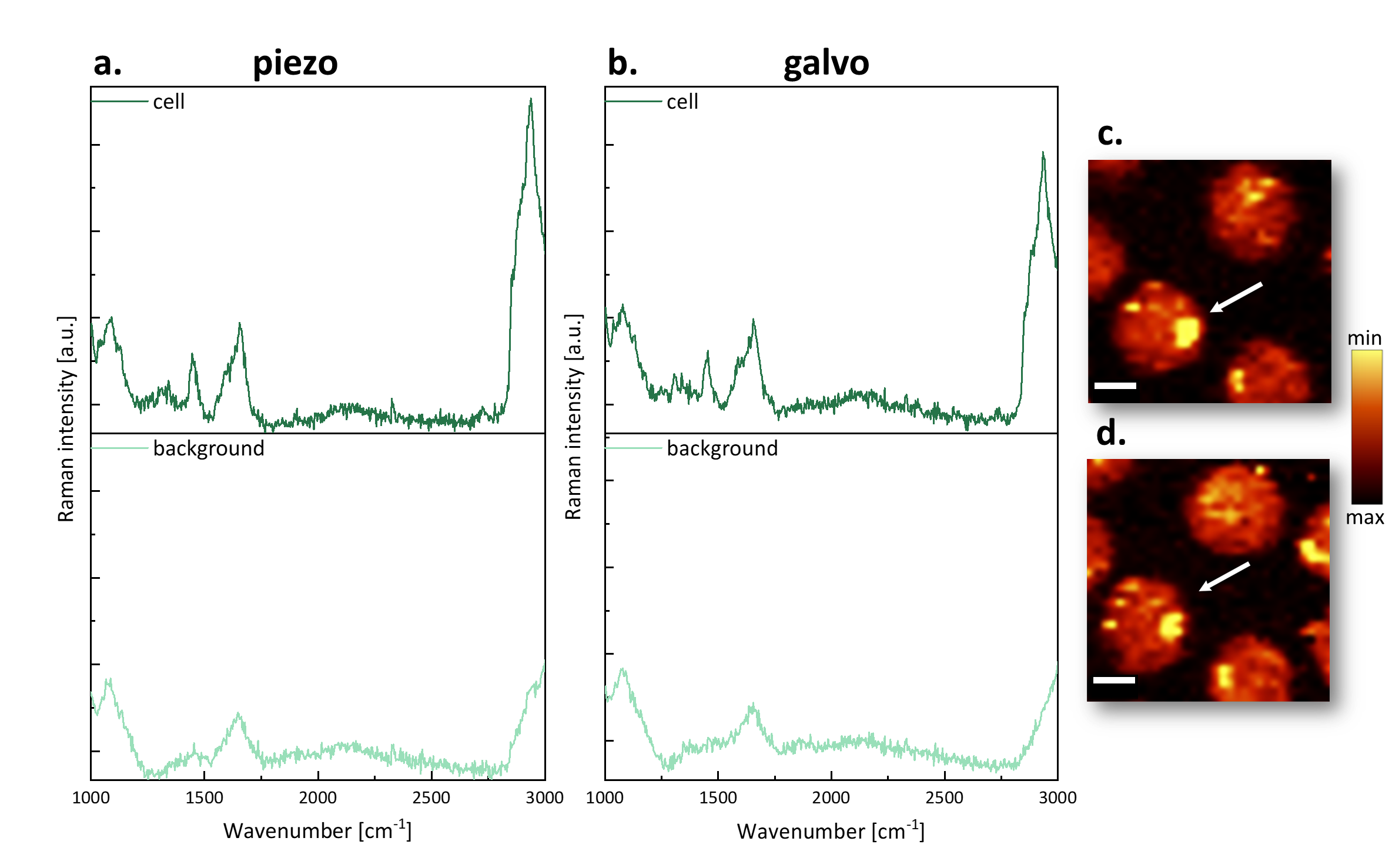

***Fig. S3*** ***Galvo-mirror vs. piezo-stage scan using RS.*** *Two scanning models were used and RS images and mean cellular spectra are shown, respectively:* ***a.*** *using piezo-scan,* ***b.*** *using galvo-scan.* ***c-d.*** *Cellular morphology presented as RS images obtained by the integration of the selected Raman bands at 2835-3025 cm^−1^ (organic matter) using piezo-scan and galvo-scan, respectively. Scale bar: 6 µm.*

When cells are measured with the TRAM system, they must be placed on glass bottom dishes to be able to perform measurements using all techniques simultaneously (RS/SRS/CARS). Unfortunately, in the case of RS imaging of such small objects as single cells (of the order of 15 µm), the influence of bands originating from the substrate (especially in the range below 1000 cm^-1^) is apparent. In the case of even smaller cells (such as lymphocytes of the order of 10 µm in size), the influence of substrate signals on the RS spectrum is even greater. This should be taken into account when planning measurements and setting the pipeline for data analysis.
